# Comparative metabonomic investigations of *Schistosoma japonicum* from SCID mice and BALB/c mice: clues to developmental abnormality of schistosome in the immunodeficient host

**DOI:** 10.1101/450858

**Authors:** Rong Liu, Wen-jun Cheng, Hong-bin Tang, Qin-ping Zhong, Zhen-ping Ming, Hui-fen Dong

## Abstract

It has been discovered that the development of schistosome is hampered in immunodeficient mice, e.g. nude mice lacking T-lymphocytes and the severe combined immune deficient (SCID) mice lacking both T- and B-lymphocytes. However, it’s still unresolved about the underlying regulatory mechanisms of the retarded growth and development of schistosomes in their immunodeficient definitive host. In this study, therefore, five replicates of male or female *Schistosoma japonicum* samples with twenty male or female worms in each sample, were collected from SCID mice or BALB/c mice at five weeks post infection and used to perform metabonomic analysis using liquid chromatography tandem mass spectrometry (LC-MS/MS) platform, for elucidating the growth and development regulation of schistosome in their definitive hosts from the metabolomic aspect. Based on the identified 1015 ion features in ESI+ mode and 342 ion features in ESI-mode, multivariate modelling methods including the Principal Component Analysis (PCA), Partial Least Squares Discriminant Analysis (PLS-DA) and Orthogonal Partial Least Squares Discriminant Analysis (OPLS-DA) identified distinct metabolic profiles that clearly differentiated both male and female worms in SCID mice from those in BALB/c mice, respectively. Common and uniquely perturbed metabolites and their involved metabolic pathways were identified in male and female worms from SCID mice when compared with those from BALB/c mice. The results also revealed that more differential metabolites were found in female worms (one metabolite was up-regulated and forty metabolites were down-regulated) than male worms (nine metabolites were up-regulated and twenty metabolites were down-regulated) between SCID mice and BALB/c mice. The top five increased metabolites of male worms in SCID mice when compared with those in BALB/c mice were PC(22:6/20:1), L-allothreonine, L-serine, glycerophosphocholine and 5-aminoimidazole ribonucleotide. And the top five decreased metabolites of male worms in SCID mice when compared with those in BALB/c mice were PC(16:0/0:0), PAF C-16, PE(18:1/0:0), adenosine and butenoyl PAF. Most of the differential metabolites of female worms in SCID mice had lower levels when compared with the normal female worms in BALB/c mice, except for retinyl ester with a higher level. The top five decreased metabolites of female worms in SCID mice when compared with those in BALB/c mice were adrenic acid, 5-phosphoribosylamine, PC(16:0/0:0), PC(22:6/20:1) and ergothioneine. The involved metabolic pathways of the differential metabolites in male worms between SCID mice and BALB/c mice mainly included taurine and hypotaurine metabolism, glycerophospholipid metabolism, sphingolipid metabolism, arachidonic acid metabolism, alpha-linolenic acid metabolism, etc. The involved metabolic pathways of differential metabolites in female worms included mainly pyrimidine metabolism, sphingolipid metabolism, arachidonic acid metabolism, glycerophospholipid metabolism, tryptophan metabolism, etc. These findings suggested a correlation between the retarded growth and development of schistosome in SCID mice and their perturbed metabolic profiles, which also provided a new insight into the regulation mechanisms of growth and development of *S. japonicum* worms from the metabolic level, and provided clues for discovery of drugs or vaccines against the parasites and parasitic disease.

**Author summary:** The growth and development of schistosome has been discovered hampered in the immunodeficient hosts. But it remains unresolved about the molecular mechanisms involved in this. In this study, we tested and compared the metabolic profiles of the male and female *Schistosoma japonicum* worms collected from SCID mice or BALB/c mice at five weeks post infection using liquid chromatography tandem mass spectrometry (LC-MS/MS) platform. There were 1015 ion features in ESI+ mode and 342 ion features in ESI-mode were identified, and distinct metabolic profiles were found to clearly differentiate both male and female worms in SCID mice from those in BALB/c mice, respectively. The results also found more differential metabolites in female worms than in male worms between SCID mice and BALB/c mice. The enriched metabolic pathways of the differential metabolites in male worms between SCID mice and BALB/c mice included taurine and hypotaurine metabolism, glycerophospholipid metabolism, sphingolipid metabolism, arachidonic acid metabolism, alpha-linolenic acid metabolism, etc. And the enriched metabolic pathways of differential metabolites in female worms included pyrimidine metabolism, sphingolipid metabolism, arachidonic acid metabolism, glycerophospholipid metabolism, tryptophan metabolism, etc. The findings in this study suggested an association between the developmentally stunted schistosome and their perturbed metabolites and metabolic pathways, which provided a new insight into the regulation mechanisms of growth and development of *S. japonicum* worms from the metabolic level, and clues for discovery of drugs or vaccines against the parasites and disease.

## Introduction

Schistosomiasis, caused by infection with a parasitic blood fluke of the genus *Schistosoma,* of which *Schistosoma mansoni, S. haematobium* and *S. japonicum* are of particular health significance, is still one of the most serious neglected tropical disease in the endemic countries [1, 2]. Unlike other trematodes, adult schistosomes are dioecious and display a fascinating codependency in that the female worm is dependent on the male to grow and sexually mature, by residing in the male’s gynecophoral canal [3]. The adult worms in pairs inhabit the mesenteric veins in the portal venous system of host, and every sexually mature female worm release thousands of eggs each day. And eggs deposited in the liver, intestinal wall and other tissues are unfortunately the main pathogenic factor to the severe schistosomiasis [3]. The highly evolved host-parasite relationship, especially that between schistosomes and their definitive hosts, is complex and long-lived [4-9].

Interestingly, however, it was found that schistosomes showed retarded growth, development and reproduction in the immunodeficient mammalian hosts, resulting in attenuated pathogenesis with decreased egg-laying and hepatic granulomas formation in the hosts [10-15]. Some researches focusing mainly on the host revealed that the host’s factors interleukin (IL)-2 and IL-7 indirectly modulated the development of blood fluke through CD4+ T cells lymphocytes [10, 11, 14]. TNF was also identified to participate in maintaining the viability of adult worms with independence of the receptors TNFR1 and TNFR2 [6]. However, few researches is available about the schistosomes’ molecular regulation on their growth and development.

Metabolic profile investigation is a promising approach to identify the key molecules or signaling pathways competent for addressing the phenotypic differences between worms from different hosts [16-29]. Ultra high performance liquid chromatography and mass spectroscopy (HPLC-MS) is capable of simultaneously detecting a wide range of small molecule metabolites and providing a “metabolic fingerprint” of biological samples, and has been used as a well-established analytical tool with successful application in different fields, e.g., studying of disease progress, detection of metabolites of inborn defects, phenotypic differentiation of experimental animal models [30-32]. In this study, therefore, we tested and compared the metabonomic perturbations of *S. japonicum* worms with sex separation from the severe combined immunodeficient (SCID) mice at the fifth week post-infection, which were compared with those from BALB/c mice as the normal control. The results will provide new insights into understanding of the molecular regulations of growth and development of schistosomes in their hosts from the metabolic level, as well as clues for discovery of drugs and vaccines against the parasites and disease.

## Materials and Methods

### Ethics statement

All experiments using the *S. japonicum* parasite, *Oncomelania hupensis* (*O. hupensis*) snails, and mice were performed under protocols approved by Wuhan University Center for Animal Experiments (WUCAE) according to the Regulations for the Administration of Affairs Concerning Experimental Animals of China (Ethical approval number: 2016025).

### Parasites and animals

*O. hupensis* snails infected with *S. japonicum* were purchased from the Institute of Parasitic Disease Control and Prevention, Hunan Province, China. Immunocompetent BALB/c mice and severe combined immunodeficient (SCID) mice of BALB/c genetic background, approximately 6∼8 weeks old, were purchased from Beijing Hua Fu Kang Bioscience Co. Inc (http://www.hfkbio.com/) via WUCAE. Cercariae were released by exposing the infected snails in aged tap water under a light for a minimum of 2 hours at 25°C, and were used to infected the above two kinds of mice via percutaneous exposure at approximately 40±1 cercariae per mouse after twelve days of acclimatization. Adult worms were collected by hepato-portal perfusion of mice with phosphate buffered solution (PBS) on the 35^th^ day post infection according to our previous research [12]. The worms were washed with PBS twice and separated manually using dissecting needle carefully under an anatomical lens if necessary, and were finally allocated 20 worms for each aliquot labeled as (1) IB-MALE, (2) IB-FEMALE, (3) IS-MALE or (4) IS-FEMALE, respectively. All the samples were frozen immediately in liquid nitrogen and then stored at -80 °C until use for metabolite extraction. In addition, the blood of mice was collected and serum was isolated and stored at -80 °C for serum metabolomics investigation, the paper about which was submitted elsewhere.

### Sample preparation

Totally 20 frozen schistosome samples in the above four groups with five replicate samples in each group were sent to Wuhan Anlong Kexun Co., LTD (www.anachro.com.cn) for metabonomic detection and analysis using HPLC-MS/MS. The LC-MS grade methanol and acetonitrile was purchased from Merck & Co., Inc., and formic acid was from Sigma-Aldrich Co. LLC. Other reagents were all analytically pure. The schistosome samples were thawed and ground after adding 0.5 ml of methanol/distilled water (8:2, v/v) with 4 μg/ml of 2-Chloro-L-phenylalanine as the internal standard substance, and were then centrifuged at 13000 rpm under 4 °C for 10 min. 200 μl of supernatant from each sample was carefully transferred to a vial of autosampler for examination. All samples were kept at 4°C and analyzed in a random manner. Additionally, isometric supernatant from each sample of the above four groups were mixed for QC sample. The QC sample was run after every 2 tested samples to monitor the stability of the system.

### Metabolomics analysis by HPLC-MS/MS

Liquid chromatography was performed on a 1290 Infinity UHPLC system (Agilent Technologies, Santa Clara, CA, U.S.A.). The separation of all samples was performed on an ACQUITY UPLC @HSS T3 column (Waters, U.K.) (100 mm * 2.1 mm, 2.5 μm). A gradient elution program was run for chromatographic separation with mobile phase A (0.1% formic acid in water) and mobile phase B (0.1% formic acid in acetonitrile) as follows: 0∼2 min, 95%A-95%A; 2∼13 min, 95%A-5%A; 13∼15 min, 5%A-5%A. The sample injection volume was 3 μL and the flow rate was set as 0.4 mL/min. The column temperature was set at 25°C, and the post time was set as 5 min.

A 6538 UHD and Accurate-Mass Q-TOF (Agilent Technologies, Santa Clara, CA, USA) equipped with an electrospray ionization (ESI) source was used for mass spectrometric detection. The electrospray ionization mass spectra for sample analysis were acquired in both positive ion mode (ESI+) and negative ion mode (ESI-). The operating parameters were as follows: capillary, 4000 V (ESI+) or 3000 V (ESI-); sampling cone: 45 V; source temperature: 110 °C (ESI+) or 120 °C (ESI-); desolvation temperature: 350 °C; desolvation gas, 11 L/min; source offset (skimmer1): 60 V; TOF acquisition mode: sensitivity (ESI+) or sensitivity (ESI-); acquisition method, continuum MSE; TOF mass range: 100-1000 Da; scan time: 0.2 s; collision energy function 2: trap CE ramp 20 to 40 eV. Quality control (QC) samples were used in order to assess the reproducibility and reliability of the LC-MS/MS system. QC samples prepared as mentioned above were used to provide a ‘mean’ profile representing all analyses encountered during the analysis. The pooled ‘QC’ samples were run before and after every two study samples to ensure system equilibration. Two reference standard compounds purine (C_5_H_4_N_4_) (with m/z 121.0509 in ESI+ mode and m/z 119.0363 in ESI-mode) and hexakis (1H,1H,3H-tetrafluoro-pentoxy)-phosphazene (C_18_H_18_O_6_N_3_P_3_F_24_) (with m/z 922.0098 in ESI+ mode and m/z 966.0007 in ESI-mode) were continuously infused into the system to allow constant mass correction during the run.

### Metabolic data analysis

Raw spectrometric data were uploaded to and analyzed with the MassHunter Qualitative Analysis B.04.00 software (Agilent Technologies, USA) for untargeted peak detection, peak alignment, peak grouping, normalization and integration on each full data set (study samples and QC samples). The molecular features, characterized by retention time (RT), chromatographic peak intensity, and accurate mass, were obtained by using the Molecular Feature Extractor algorithm. The features were then analyzed with the MassHunter Mass Profiler Professional software (Agilent Technologies). Only features with an intensity of ≥ 20,000 counts (approximately three times the detection limit of the LC-MS/MS instrument used in this study) that were found in at least 80% of the samples at the same sampling time point were kept for further processing. Next, a tolerance window of 0.15 min and 2 mDa was used for alignment of retention time and m/z values, and the data were also normalized by the internal standard 2-Chloro-L-phenylalanine added when sample preparation. The data matrix was then mean-centered and Pareto-scaled prior to multivariate analysis (MVA) using Principal Component Analysis (PCA), Partial Least Squares Discriminant Analysis (PLS-DA) and Orthogonal Partial Least Squares Discriminant Analysis (OPLS-DA) to discriminate comparison groups using the function module *Statistical Analysis* on the online application *MetaboAnalyst* (http://www.metaboanalyst.ca/) [33-35]. The quality of the models was evaluated with the relevant parameters R^2^ and Q^2^, which were discussed elsewhere (Lee et al., 2003). And differences between groups (IS-MALE *vs.* IB-MALE, and IS-FEMALE *vs.* IB-FEMALE) were determined using Student’s *t*-test, and the adjusted *P* value (false discovery rate, FDR) of <0.05 was considered to be of statistical significance. Fold change (FC) analysis, which was used to show how the selected differential metabolites varied between the compared groups, was also performed to further filter the features/metabolites of particular concern with an FC of ≥1.2 or ≤0.8 between the compared groups.

The structure identification of the differential metabolites was based on the methods described as follows. Briefly, the element compositions of the metabolites were first calculated with MassHunter software from Agilent based on the exact mass, the nitrogen rule, and the isotope pattern. Then, the elemental composition and exact mass were used for open source database searching, including LIPIDMAPS (http://www.lipidmaps.org/), HMDB (http://www.hmdb.ca/), METLIN (http://metlin.scripps.edu/), and MassBank (http://www.massbank.jp/). Next, MS/MS experiments were performed to obtain structural information via the interpretation of the fragmentation pattern of the metabolite. The MS/MS spectra of possible metabolite candidates in the databases were also searched and matched.

*MetaboAnalyst* was used to perform metabolic pathway analysis of the differentially expressed metabolites. The identified pathways associated with the abnormal growth and development of schistosome in SCID mice are presented according to the *P*-values from the pathway enrichment analysis (y-axis) and pathway impact values from pathway topology analysis (x-axis), with the most impacted pathways colored in red color.

## Results

### Metabolic profiles

All total ion chromatograms (TIC) of QC samples exhibited stable retention times without obvious peaks’ drifts (Figure S1), which indicated good capability of the LC-MS/MS based-metabolomics approach used in this study. Totally, 1015 ion features in ESI+ mode and 342 ion features in ESI-mode were obtained in all the male or female *S. japonicum* worms samples, respectively. The stability and reproducibility of the HPLC-MS/MS method was evaluated by performing principal components analysis (PCA) on all the samples, together with 10 QC samples. The QC samples are generally clustered closely to each other and are separated from the tested samples in the two-dimensional PCA score plots (Figure 1-A, B) and PLS-DA score plots (Figure 1-C, D), though a moderate separation among the QC samples in ESI+ mode was observed (Figure 1-A), which confirms good stability and reproducibility of the chromatographic separation during the whole sequence. In addition, although the male worms (both IS-MALE and IB-MALE) were clearly separated from the female worms (both IS-FEMALE and IB-FEMALE), the male worms IS-MALE and IB-MALE were partially overlapped in the two-dimensional PCA score plots in both ion modes (Figure 1-A, B), while the female worms IS-FEMALE and IB-FEMALE were completely separated in ESI-mode (Figure 1-B), which indicated larger differences between male and female worms than the differences between the worms of the same sex derived from two different hosts, and MALE. Similar results were also found in the two-dimensional PLS-DA models performed on all the samples, and it yielded distinct separation of the tested four groups of worms in both ESI+ (Figure 1-C) and ESI-mode (Figure 1-D).

**Figure 1.**
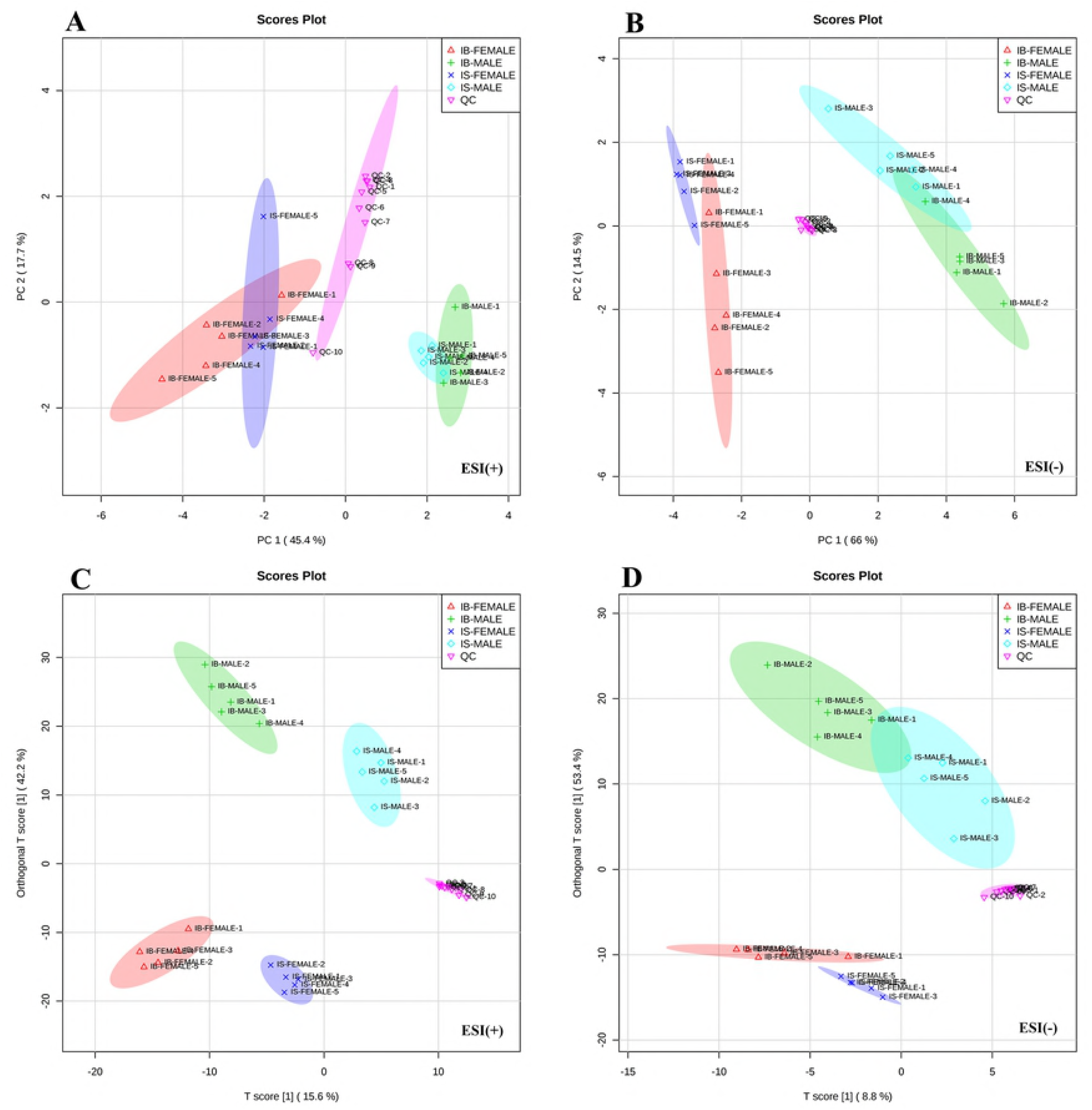
Differential metabolic profiles of male and female *S. japonicum* worms between SCID mice and BALB/c mice at 35 days post infection. Principal component analysis (PCA) score scatter plots of metabolites obtained from LC-MS/MS fingerprints in ESI+ (A) and ESI-mode (B). Partial least-squares discriminant analysis (PLS-DA) separating metabolites of the four groups of worms and QC sample in ESI+ (C) and ESI-mode (D). IB-MALE denotes the male worms from BALB/c mice, which are marked with green plus sign. IB-FEMALE denotes the female worms from BALB/c mice, which are marked with red triangles. IS-MALE denotes the male worms from SCID mice, which are marked with blue diamonds. IS-FEMALE denotes the female worms from SCID mice, which are marked with purple cross. QC denotes the quality control samples, which are marked with pink triangles.

### Metabolomic profiles distinguish between *S. japonicum* worms from SCID mice and those from BALB/c mice

Score scatter plots for two-dimensional OPLS-DA model in both ESI+ mode (Figure 2-A) and ESI-mode (Figure 2-B) showed good discrimination between the male worms from SCID mice and those from BALB/c mice (IS-MALE *vs.* IB-MALE), which was also demonstrated in the heatmap based on the differential metabolites of the 10 male worms samples (Figure 2-C). Likewise, the score plots for two dimensional OPLS-DA mode in both ESI+ mode (Figure 2-D) and ESI-mode (Figure 2-E) also showed distinct group separation between the female worms samples from SCID mice and those from BALB/c mice (IS-FEMALE *vs.* IB-FEMALE), which was further supported by the heatmap constructed based on the 10 female worms samples (Figure 2-F).

**Figure 2.**
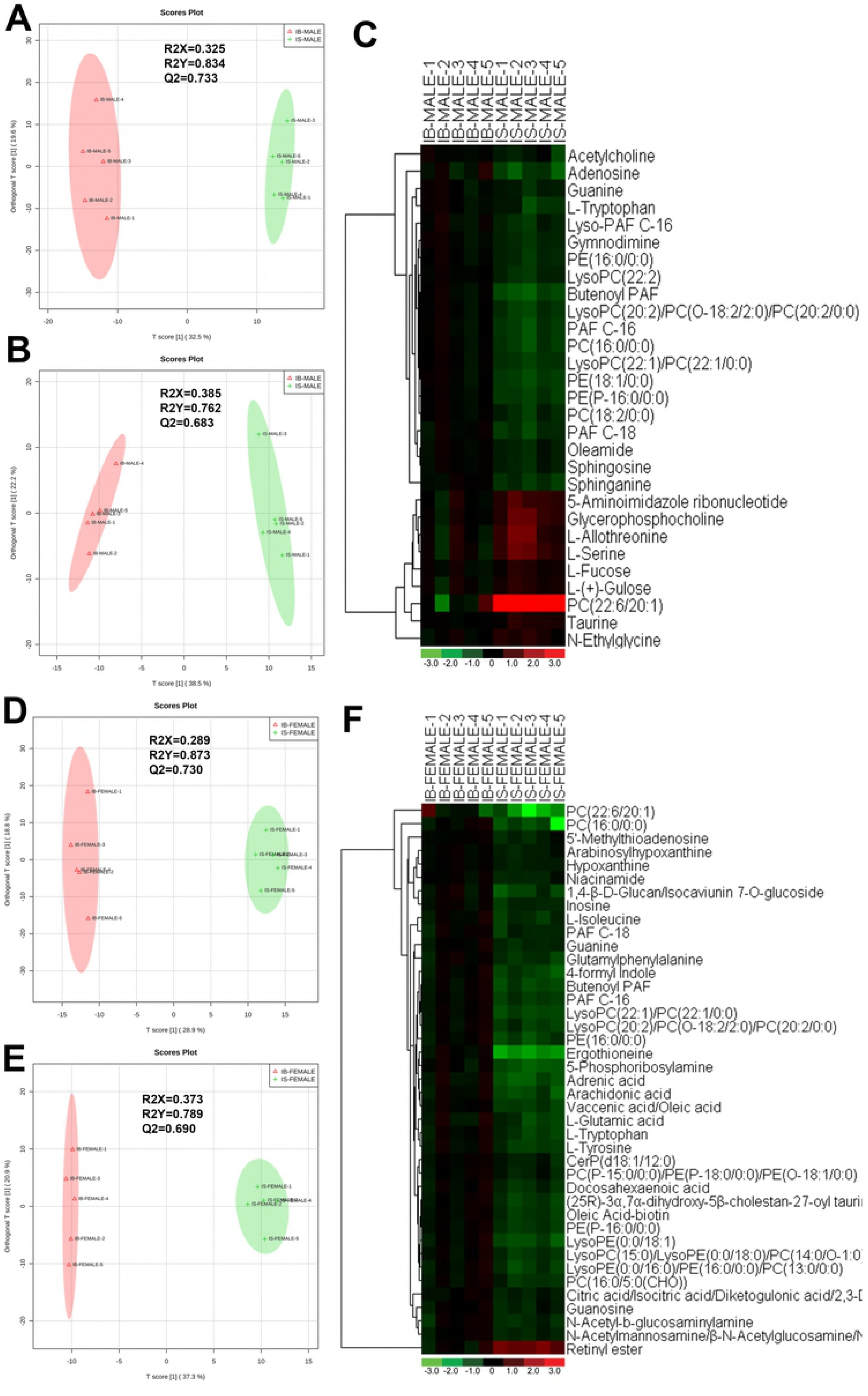
Discrimination between the *S. japonicum* worms from SCID mice and those from BALB/c mice with *S. japonicum* infection for 35 days based on ESI+ and ESI-mode-derived metabolic phenotypes and heatmaps of the differential metabolites between the compared groups. **A-B**: Orthogonal partial least-squares discriminant analysis (OPLS-DA) score plots in ESI+ mode (A) and ESI-mode (B) for comparison between male worms from SCID mice and those from BALB/c mice (IB-MALE denotes the male worms from BALB/c mice and IS-MALE denotes the male worms from SCID mice). **C**: Heatmap of the differential metabolites between male worms from SCID mice and those from BALB/c mice. **D-E**: Orthogonal partial least-squares discriminant analysis (OPLS-DA) score plots in ESI+ mode (A) and ESI-mode (B) for comparison between female worms from SCID mice and those from BALB/c mice (IB-FEMALE denotes the female worms from BALB/c mice and IS-FEMALE denotes the female worms from SCID mice). **F:** Heatmap of the differential metabolites between female worms from SCID mice and those from BALB/c mice. For the heatmaps, normalized signal intensities (log2 transformed and row adjustment) are visualized as a color spectrum and the scale from least abundant to highest ranges is from -3.0 to 3.0 as shown in the colorbar. Green indicates decreased expression, whereas red indicates increased expression of the detected metabolites between compared groups.

### Patterns of metabolites with differential amount in schistosome

Twenty-nine differential ion features/metabolites (FDR<0.05, and FC ≥1.2 or ≤0.8), with nine increased and twenty decreased, were identified between IS-MALE *vs.* IB-MALE (Table S1, Figure 2-C). Five of the increased metabolites, PC(22:6/20:1) (which was traditionally named as lecithin), L-allothreonine, L-serine, glycerophosphocholine, and 5-aminoimidazole ribonucleotide, even had an FC >1.5, particularly for PC(22:6/20:1) involved in the of the twenty decreased metabolites had an FC <0.5 between IS-MALE *vs.* IB-MALE. Meanwhile, forty-one differential features/metabolites were identified between IS-FEMALE and IB-FEMALE (Table S2, Figure 2-F). Retinyl ester, a metabolite of the retinol metabolism pathway, is the only metabolite that showed up-regulated between IS-FEMALE and IB-FEMALE. Four of the remained forty decreased metabolites, 5-phosphoribosylamine, PC(16:0/0:0), PC(22:6/20:1) and ergothioneine, even had an FC <0.5 between IS-FEMALE and IB-FEMALE.

Comparison of differential metabolic profiles between IS-MALE *vs*. IB-MALE and IS-FEMALE *vs.* IB-FEMALE found that eleven features/metabolites were common in their differential metabolites (Table 1, Figure 3-A, B). Most of the common differential metabolites in both male and female worms had the similar decrease trend in the SCID mice, except that PC(22:6/20:1) increased in male worms but decreased in female worms in SCID mice compared with BALB/c mice (Table 1 and Figure 3-B). After removing the common differential metabolites, eighteen differential metabolites were distinct in IS-MALE *vs.* IB-MALE (Table 2, Figure 3-C), and thirty differential metabolites were distinct in IS-FEMALE *vs.* IB-FEMALE (Table 3, Figure 3-D), which is more than male worms. These differential metabolites common and distinct in male worms or female worms in SCID mice compared with BALB/c mic may be associated with the abnormal growth and development of worms in SCID mice, and the differential metabolites distinct in male worms or female worms should be associated with larger differences between IS-FEMALE and IB-FEMALE than those between IS-MALE and IB-MALE.

**Figure 3.**
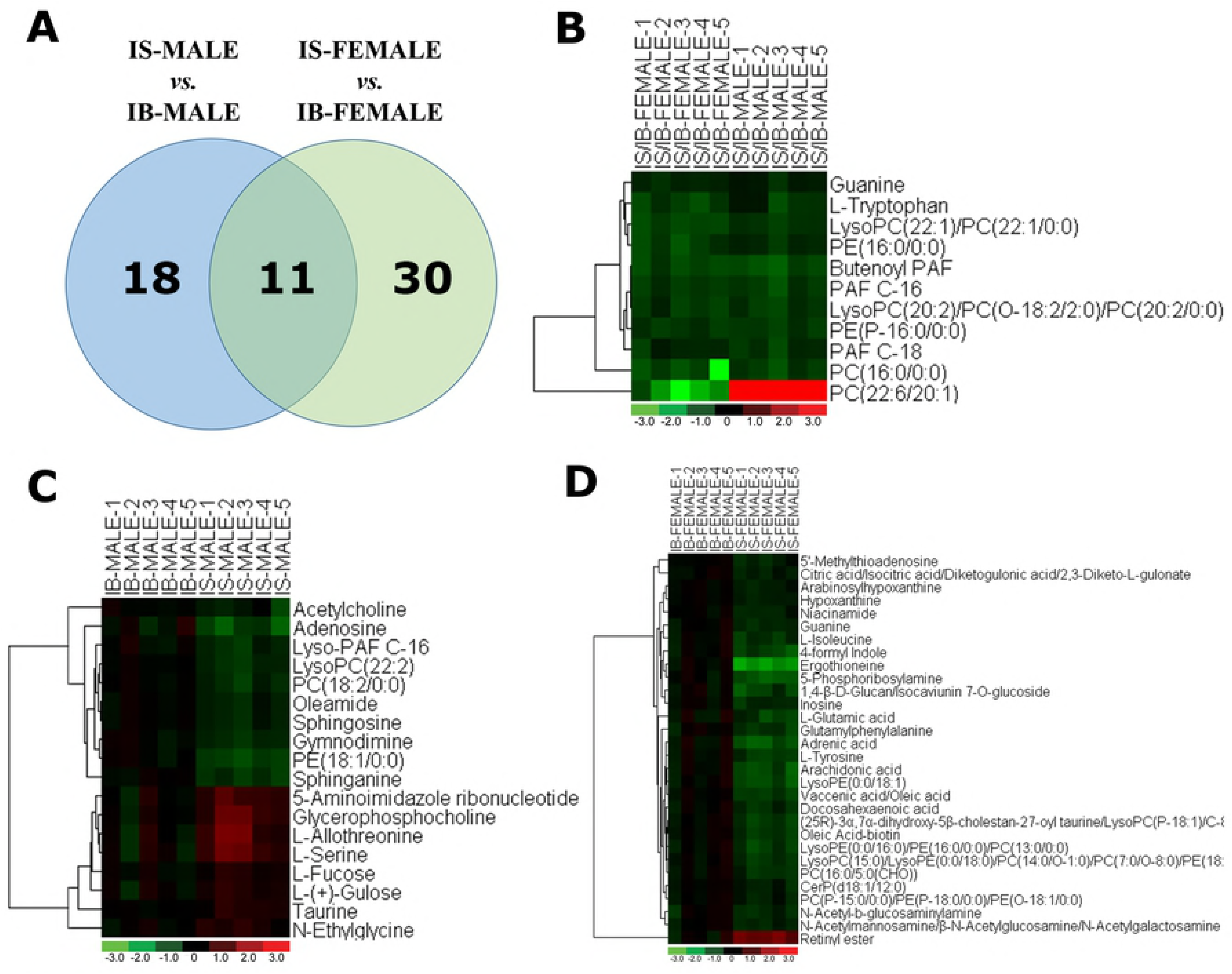
Comparison of metabolomics between the *S. japonicum* worms from SCID mice and those from BALB/c mice with *S. japonicum* infection for 35 days heatmaps of their differential metabolites. **A:** Differential metabolites across comparison groups showing unique and common metabolites between IS-MALE *vs.* IB-MALE and IS-FEMALE *vs.* IB-FEMALE. Venn diagram displays comparatively the differentially expressed metabolites. All the differentially expressed metabolites are clustered into two comparison groups represented by two circles. The sum of all the figures in one circle represents the number of differentially expressed metabolites in one comparison group (e.g. IS-MALE *vs.* IB-MALE). The overlapping part of the two circles represents the number of differentially expressed metabolites shared between the two comparison groups. The single-layer part represents the number of metabolites distinctly found in a certain comparison group. **B-C:** Heatmap of the common (B) and distinct (C is for male worms and D is for female worms) differential metabolites between male worms and female worms. For the heatmaps, normalized signal intensities (log2 transformed and row adjustment) are visualized as a color spectrum and the scale from least abundant to highest ranges is from -3.0 to 3.0 as shown in the colorbar. Green indicates decreased expression, whereas red indicates increased expression of the detected metabolites between compared groups.

**Table 1.**
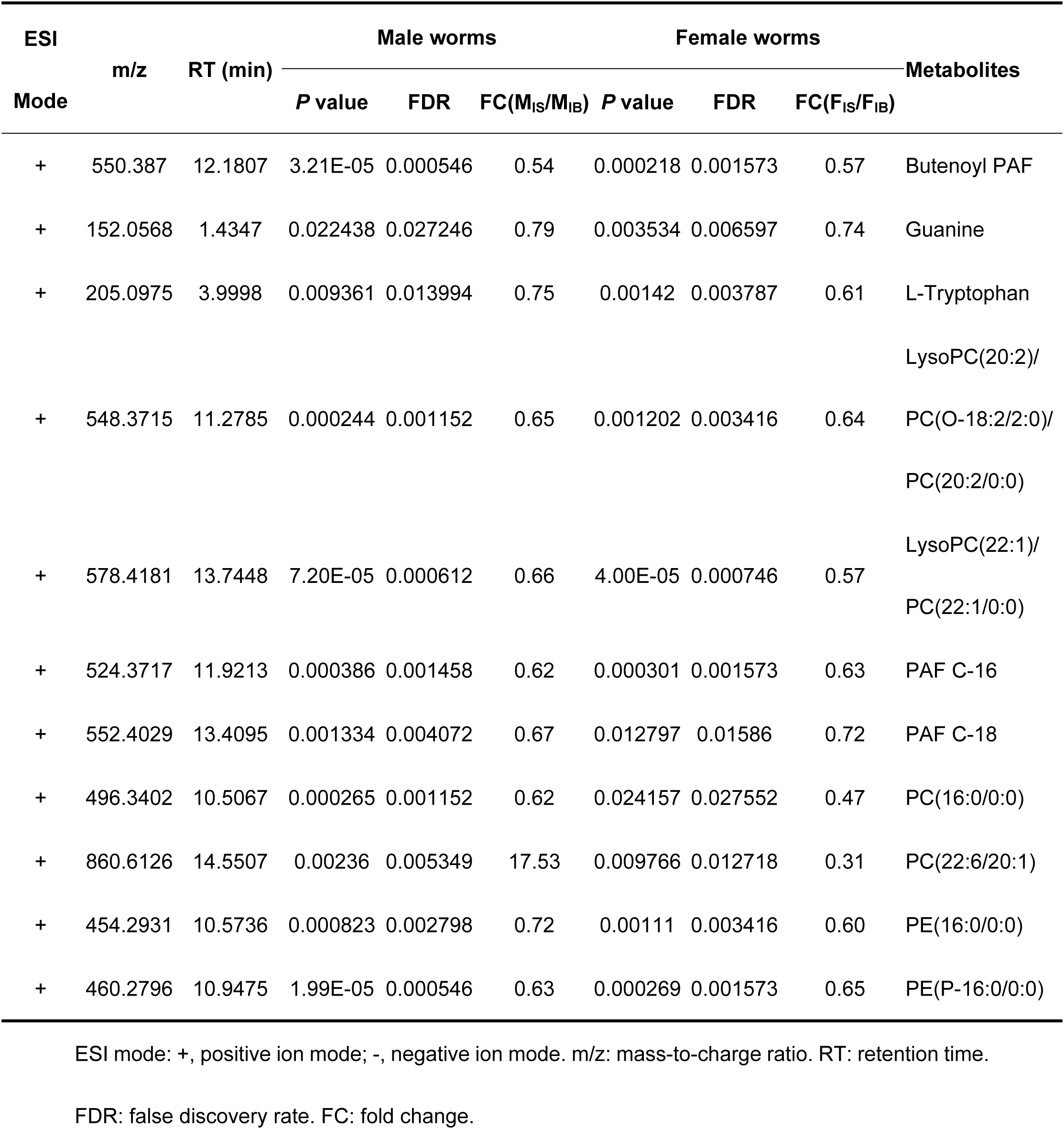
List of the common differential metabolites in male and female worms from SCID mice

**Table 2.**
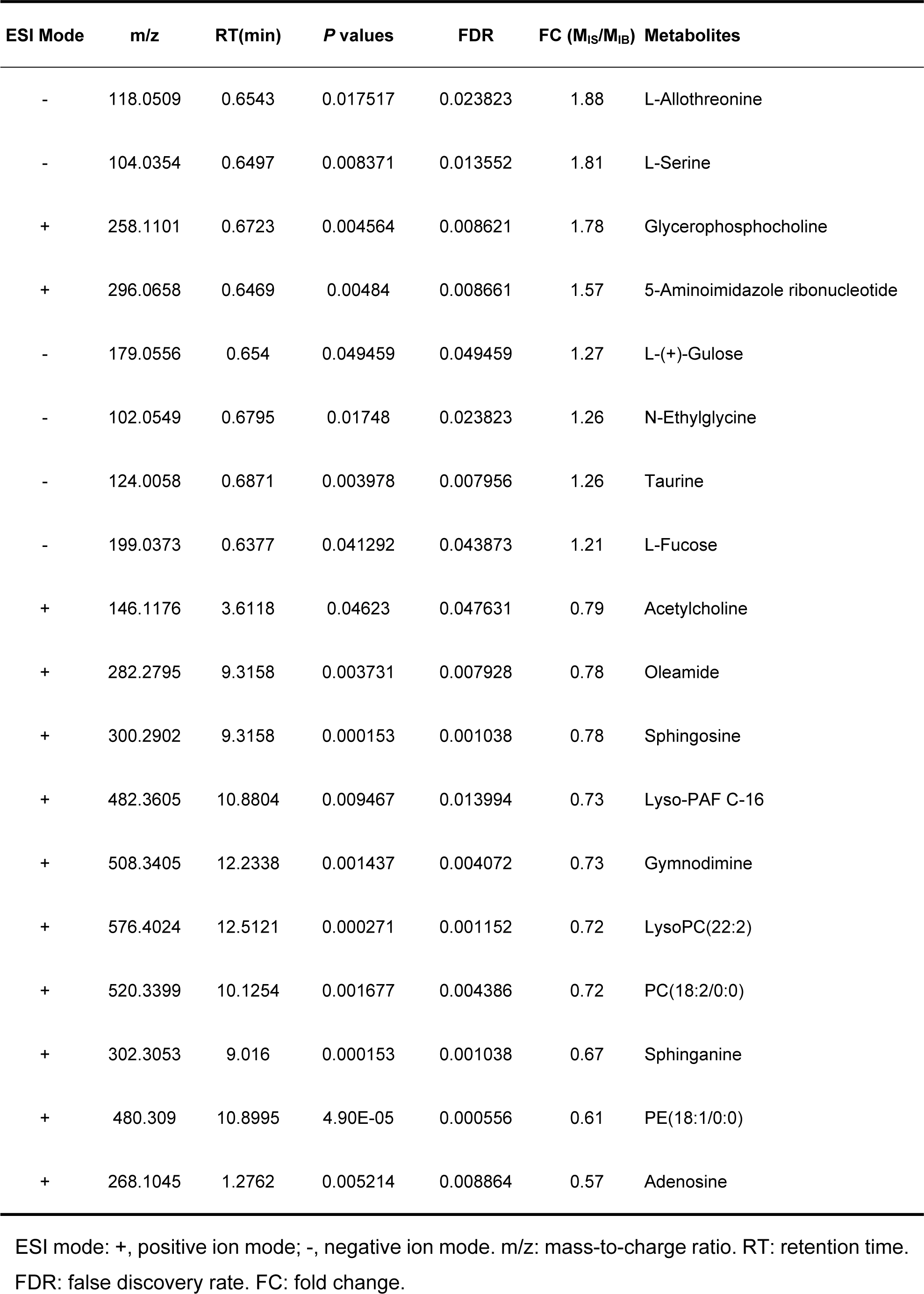
List of the male worm-specific differential metabolites between SCID mice and BALB/c mice

| ESI Mode | m/z | RT(min) | P values | FDR | FC (M<sub>IS</sub>/M<sub>IB</sub>) | Metabolites |
| --- | --- | --- | --- | --- | --- | --- |
| - | 118.0509 | 0.6543 | 0.017517 | 0.023823 | 1.88 | L-Allothreonine |
| - | 104.0354 | 0.6497 | 0.008371 | 0.013552 | 1.81 | L-Serine |
| + | 258.1101 | 0.6723 | 0.004564 | 0.008621 | 1.78 | Glycerophosphocholine |
| + | 296.0658 | 0.6469 | 0.00484 | 0.008661 | 1.57 | 5-Aminoimidazole ribonucleotide |
| - | 179.0556 | 0.654 | 0.049459 | 0.049459 | 1.27 | L-(+)-Gulose |
| - | 102.0549 | 0.6795 | 0.01748 | 0.023823 | 1.26 | N-Ethylglycine |
| - | 124.0058 | 0.6871 | 0.003978 | 0.007956 | 1.26 | Taurine |
| - | 199.0373 | 0.6377 | 0.041292 | 0.043873 | 1.21 | L-Fucose |
| + | 146.1176 | 3.6118 | 0.04623 | 0.047631 | 0.79 | Acetylcholine |
| + | 282.2795 | 9.3158 | 0.003731 | 0.007928 | 0.78 | Oleamide |
| + | 300.2902 | 9.3158 | 0.000153 | 0.001038 | 0.78 | Sphingosine |
| + | 482.3605 | 10.8804 | 0.009467 | 0.013994 | 0.73 | Lyso-PAF C-16 |
| + | 508.3405 | 12.2338 | 0.001437 | 0.004072 | 0.73 | Gymnodimine |
| + | 576.4024 | 12.5121 | 0.000271 | 0.001152 | 0.72 | LysoPC(22:2) |
| + | 520.3399 | 10.1254 | 0.001677 | 0.004386 | 0.72 | PC(18:2/0:0) |
| + | 302.3053 | 9.016 | 0.000153 | 0.001038 | 0.67 | Sphinganine |
| + | 480.309 | 10.8995 | 4.90E-05 | 0.000556 | 0.61 | PE(18:1/0:0) |
| + | 268.1045 | 1.2762 | 0.005214 | 0.008864 | 0.57 | Adenosine |

3 ESI mode: +, positive ion mode; -, negative ion mode. m/z: mass-to-charge ratio. RT: retention time.
FDR: false discovery rate. FC: fold change.

**Table 3.** List of the female worm-specific differential metabolites between SCID mice and BALB/c mice

| ESI Mode | m/z | RT(min) | P values | FDR | FC (F<sub>IS</sub>/F<sub>IB</sub>) | Metabolites |
| --- | --- | --- | --- | --- | --- | --- |
| - | 301.2171 | 12.9477 | 0.000132 | 0.001233 | 2.34 | Retinyl ester |
| + | 269.0885 | 1.4201 | 0.003199 | 0.006398 | 0.80 | Arabinosylhypoxanthine |
| - | 464.3143 | 12.3082 | 0.025349 | 0.027741 | 0.80 | PC(P-15:0/0:0)/PE(P-18:0/0:0)/PE(O-18:1/0:0) |
| + | 137.046 | 1.4199 | 0.000225 | 0.001573 | 0.79 | Hypoxanthine |
| - | 191.0178 | 0.8004 | 0.008998 | 0.011998 | 0.79 | Citric acid/Isocitric acid/Diketogulonic acid/<br>2,3-Diketo-L-gulonate |
| - | 596.3926 | 13.4124 | 0.010324 | 0.013139 | 0.79 | CerP(18:1/12:0) |
| + | 284.0993 | 1.4349 | 0.025797 | 0.027741 | 0.78 | Guanosine |
| + | 298.0974 | 4.0607 | 0.00461 | 0.007822 | 0.78 | 5'-Methylthioadenosine |
| + | 123.0554 | 1.0077 | 0.001965 | 0.004584 | 0.77 | Niacinamide |
| - | 267.0717 | 1.4171 | 0.002114 | 0.004735 | 0.76 | Inosine |
| - | 256.0576 | 0.6944 | 0.007146 | 0.010815 | 0.76 | N-Acetylmannosamine/N-Acetyl-b-D-galactosamine/Beta-N-Acetylglucosamine/N-Acetylgalactosamine |
| - | 219.0968 | 0.6785 | 0.0246 | 0.027552 | 0.72 | N-Acetyl-b-glucosaminylamine |
| - | 327.2327 | 13.3711 | 0.00792 | 0.011671 | 0.71 | Docosahexaenoic acid |
| - | 592.3605 | 11.2786 | 0.004009 | 0.007241 | 0.69 | PC(16:0/5:0(CHO)) |
| + | 132.1019 | 1.172 | 0.004177 | 0.00731 | 0.68 | L-Isoleucine |
| - | 480.309 | 11.876 | 0.00122 | 0.003416 | 0.67 | LysoPC(15:0)/LysoPE(0:0/18:0)/PC(14:0/O-1:0)/PC(7:0/O-8:0)/PE(18:0/0:0)/PC(15:0/0:0) |
| - | 293.1175 | 4.4634 | 0.003346 | 0.006462 | 0.67 | Glutamylphenylalanine |
| - | 557.3195 | 10.6175 | 0.000432 | 0.001862 | 0.66 | Oleic Acid-biotin |
| + | 182.0811 | 1.1304 | 0.001879 | 0.004574 | 0.65 | L-Tyrosine |
| - | 146.0449 | 0.6795 | 0.013028 | 0.01586 | 0.64 | L-Glutamic acid |
| - | 540.3297 | 10.6169 | 0.000736 | 0.002944 | 0.64 | (25R)-3alpha,7alpha-dihydroxy-5beta-cholestan-27-oyl taurine /<br>LysoPC(P-18:1)/C-8 Ceramide-1-phosphate |
| - | 452.2769 | 10.5719 | 0.002999 | 0.006219 | 0.63 | LysoPE(0:0/16:0)/PE(16:0/0:0)/PC(13:0/0:0) |
| - | 281.2481 | 14.5325 | 7.09E-05 | 0.000794 | 0.63 | Vaccenic acid/Oleic acid |
| - | 535.1528 | 1.4173 | 0.001719 | 0.004375 | 0.59 | 1,4-beta-D-Glucan/Isocaviunin 7-O-glucoside |
| + | 146.0602 | 4.0002 | 0.000858 | 0.003004 | 0.58 | 4-formyl Indole |
| - | 303.2326 | 13.5603 | 0.000836 | 0.003004 | 0.56 | Arachidonic acid |
| - | 478.2923 | 10.9031 | 0.000403 | 0.001862 | 0.55 | LysoPE(0:0/18:1) |
| - | 331.2635 | 14.2904 | 0.000309 | 0.001573 | 0.51 | Adrenic acid |
| - | 264.005 | 0.7038 | 2.66E-05 | 0.000745 | 0.50 | 5-Phosphoribosylamine |
| + | 230.0967 | 0.7226 | 7.35E-07 | 4.12E-05 | 0.29 | Ergothioneine |

1 ESI mode: +, positive ion mode; -, negative ion mode. m/z: mass-to-charge ratio. RT: retention time.
FDR: false discovery rate. FC: fold change.

By searching against “The Human Metabolome Database” (HMDB, access via http://www.hmdb.ca/) for metabolite classification, “glycerophospholipids”, “organonitrogen compounds” and “carboxylic acids and derivatives” were found as the top three categories (≥ 3 differential metabolites involved) of differential metabolites between IS-MALE *vs.* IB-MALE (Figure 4-A). Metabolite set enrichment analysis (MSEA, access via http://www.metaboanalyst.ca/) found that “bile acid biosynthesis”, “taurine and hypotaurine metabolism”, “sphingolipid metabolism”, “retinol metabolism”, “purine metabolism”, “fructose and mannose degradation”, “ammonia recycling”, “glycine and serine metabolism”, “homocysteine degradation”, “phosphatidylethanolamine biosynthesis”, “methionine metabolism” and “selenoamino acid metabolism” were the prominently enriched metabolite sets (with adjusted *P* values < 0.05) based on the differential metabolites between IS-MALE *vs.* IB-MALE (Table S3, Figure 4-B). Meanwhile, “glycerophospholipids”, “carboxylic acids and derivatives”, “organonitrogen compounds”, “fatty acyls” and “purine nucleosides” were the top five enriched categories of differential metabolites between IS-FEMALE *vs.* IB-FEMALE (Figure 4-C). And MSEA based on the differential metabolites between IS-FEMALE *vs.* IB-FEMALE found that “retinol metabolism”, “alpha linolenic acid and linoleic acid metabolism”, “purine metabolism”, “sphingolipid metabolism” and “glutamate metabolism” were the prominently enriched metabolite sets with raw P values < 0.05 but only “retinol metabolism” has an adjusted *P* value < 0.05 (Table S4, Figure 4-D). Moreover, most (9/11) of the common differential metabolites between IS-MALE *vs.* IB-MALE and IS-FEMALE *vs.* IB-FEMALE belong to glycerophospholipids (Figure 4-E), which was enriched by MSEA to phospholipid biosynthesis (Table S5, Figure 4-F). The differential metabolites distinct in IS-MALE *vs.* IB-MALE were classified prominently into “organonitrogen compounds”, “glycerophospholipids” and “carboxylic acids and derivatives” (Figure 4-G), which were enriched prominently to “sphingolipid metabolism”, “purine Metabolism”, “methionine metabolism”, “selenoamino acid metabolism”, “bile acid biosynthesis”, “taurine and hypotaurine metabolism”, “retinol metabolism” and “betaine metabolism” (Table S6, Figure 4-H). The differential metabolites distinct in IS-FEMALE *vs.* IB-FEMALE were classified prominently into “glycerophospholipids”, “carboxylic acids and derivatives”, “fatty acyls”, “organonitrogen compounds” and “purine nucleosides” (Figure 4-I), which were enriched prominently to “retinol metabolism”, “purine metabolism”, “glutamate metabolism”, “alpha linolenic acid and linoleic acid metabolism” and “warburg effect” (Table S7, Figure 4-J).

**Figure 4.**
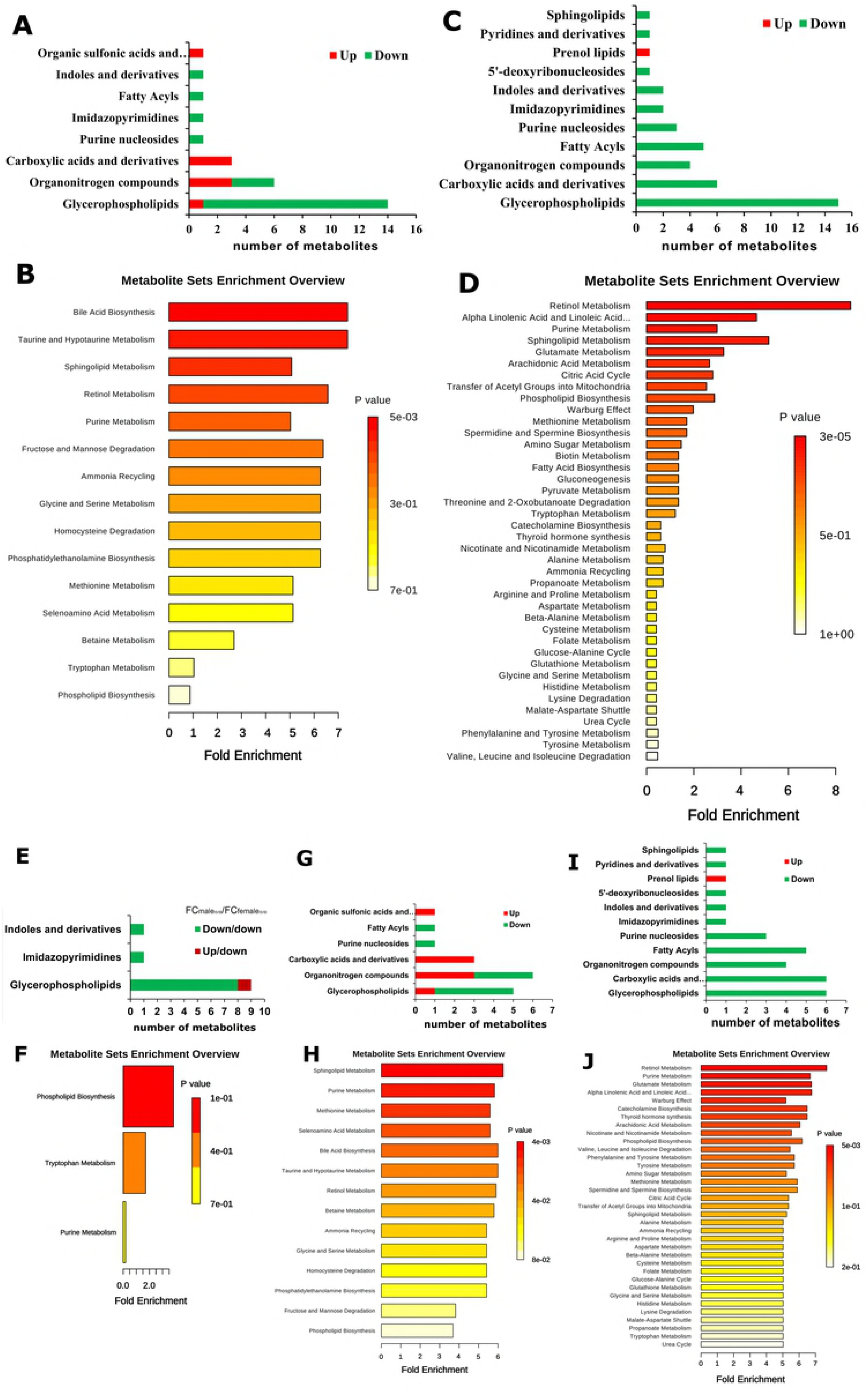
Enrichment analysis of the differential metabolites across comparison groups. **A**: Top 8 enriched metabolite terms of the differentially expressed metabolites of IS-MALE *vs.* IB-MALE. The bars on x-axis represent the number of metabolites for the chemical classes mentioned on the y-axis. **B**: Enriched metabolite sets of the differentially expressed metabolites of IS-MALE *vs.* IB-MALE. **C**: Top 11 enriched metabolite terms of the differentially expressed metabolites of IS-FEMALE *vs.* IB-FEMALE. **D**: Enriched metabolite sets of the differentially expressed metabolites of IS-FEMALE *vs.* IB-FEMALE. **E**: Enriched metabolite terms of the common differentially expressed metabolites between IS-MALE *vs.* IB-MALE and IS-FEMALE *vs.* IB-FEMALE. **F**: Enriched metabolite sets of the differentially expressed metabolites between IS-MALE *vs.* IB-MALE and IS-FEMALE *vs.* IB-FEMALE. **G**: Enriched metabolite terms of the differentially expressed metabolites distinct in IS-MALE *vs.* IB-MALE. **H**: Enriched metabolite sets of the differentially expressed metabolites distinct in IS-MALE *vs.* IB-MALE. **I**: Enriched metabolite terms of the differentially expressed metabolites distinct in IS-FEMALE *vs.* IB-FEMALE. **J**: Enriched metabolite sets of the differentially expressed metabolites distinct in IS-FEMALE *vs.* IB-FEMALE.

### Altered metabolic pathways and their biological significance

The pathway analysis performed using *MetaboAnalyst* for the involved biological pathways and biological roles of the above differentially expressed metabolites determined that the perturbed metabolic pathways reporting lower p-values and higher pathway impact in male worms from SCID mice compared with those from BALB/c mice mainly included arachidonic acid metabolism, alpha-linolenic acid metabolism, taurine and hypotaurine metabolism, sphingolipid metabolism, glycerophospholipid metabolism, and etc (Table S8, Figure S2-A). Meanwhile, more affected metabolic pathways in female worms from SCID mice compared with those from BALB/c mice were determined, and the top 5 metabolic pathways included biotin metabolism, tryptophan metabolism, purine metabolism, glyoxylate and dicarboxylate metabolism, tyrosine metabolism (Table S9, Figure S2-B).

Metabolic pathways analysis based on the common differential metabolites between IS-MALE *vs.* IB-MALE and IS-FEMALE *vs.* IB-FEMALE found that tryptophan metabolism, aminoacyl-tRNA biosynthesis, purine metabolism, glycerophospholipid metabolism, arachidonic acid metabolism and alpha-linolenic acid metabolism were commonly perturbed in both male and female worms from SCID mice compared with BALB/c mice (Table S10, Figure S2-C). Metabolic pathways analysis based on the differential metabolites distinct in IS-MALE *vs.* IB-MALE found their involved metabolic pathways included sphingolipid metabolism, glycerophospholipid metabolism, taurine and hypotaurine metabolism, purine metabolism, glycine/serine/threonine metabolism, cysteine and methionine metabolism, cyanoamino acid metabolism, glyoxylate and dicarboxylate metabolism and aminoacyl-tRNA biosynthesis (Table S11, Figure S2-D). More metabolic pathways based on the differential metabolites distinct in IS-FEMALE *vs.* IB-FEMALE were found and the top five metabolic pathways were arachidonic acid metabolism, glycerophospholipid metabolism, glycosylphosphatidylinositol (GPI)-anchor biosynthesis, alpha-linolenic acid metabolism and glyoxylate and dicarboxylate metabolism (Table S12, Figure S2-E).

## Discussion

The schistosomes exhibit dioecy and have a complex life cycle transforming between their definitive hosts - mammals and their intermediate hosts - snails. Every mature female worm lay hundreds to thousands of eggs per day, which is a complex metabolic process such as the fatty acid oxidation, etc [36]. A certain part of the released eggs are deposited in the liver, intestinal wall and other tissues, which are the key pathogenic factor to severe schistosomiasis. So, it is a feasible approach to control schistosomiasis by interfering with the growth, development and oviposition of schistosomes in their hosts. It has been already discovered that schistosome showed retarded growth, development and reproduction in the immunodeficient mammalian hosts, and exerted attenuated pathogenesis with decreased egg-laying and granulomas formation in the hosts. However, very limited knowledge about the molecular mechanism behind the distinct phenotypic abnormalities of schistosomes in their immunodeficient hosts is accessible. For the abnormal schistosomes in immunodeficient hosts, e.g. SCID mice, we believe it is a perfect model by comparative omics to identify molecules potentially participate in the regulation of growth and development of schistosomes when compared with those with normal growth and development in immunocompetent mice, e.g. BALB/c mice. In this study, therefore, an untargeted LC-MS/MS-based high-resolution metabolomic investigation was performed and distinct bio-signatures in the metabolic profiles of male and female *S. japonicum* worms in SCID mice were found when compared with those in BALB/c mice, respectively.

In the results, multivariate analysis by both PCA and PLS-DA found larger differences between IS-FEMALE and IB-FEMALE than that between IS-MALE and IB-MALE. This indicates the female schistosome worms were affected more severely than the male worms in SCID mice, which was verified by the subsequent finding that more differential metabolites were acquired in IS-FEMALE *vs.* IB-FEMALE than IS-MALE *vs.* IB-MALE. This is expectable and reasonable as the growth and development of female worms were affected by male worms as well as the host’s factors, i.e. the sexual maturation of female worms depends on pairing with male worms. In the list of differential metabolites of IS-FEMALE *vs.* IB-FEMALE, retinyl ester was the only up-regulated metabolite of female worms from SCID mice, which was enriched in “retinol metabolism”. Numerous researches reported that the retinol metabolism, in which retinyl ester is involved, regulates gametogenesis and reproduction by the product transcriptionally active retinoic acid [37-41]. In the retinol metabolism pathway of animals, all-trans retinyl esters in the body is formed by transferring a fatty acyl moiety from the sn-1 position of membrane phosphatidyl choline (e.g. PC(22:6/20:1), traditionally named as lecithin) to all-trans-retinol under the catalysis of lecithin:retinol acyltransferase (LRAT), whose orthologue in *Schistosoma* is diacylglycerol O-acyltransferase (DGAT). Unesterified all-trans-retinol, which could be reversibly liberated from all-trans retinyl esters stores through the action of a retinyl ester hydrolase (REH), is oxidized by one retinol dehydrogenase (RDH) to all-trans-retinal, which can be also reversibly transformed to all-trans-retinol by the catalysis of retinal reductase (RALR). All-trans-retinal, which is originally derived from the decomposition of proretinoid carotenoids such as dietary β-carotene, is then irreversibly oxidized by one retinal dehydrogenase (RALDH) to form transcriptionally active all-trans retinoic acid [42]. LRAT is a key enzyme involved in retinoids homeostasis and is regulated in response to retinoic acid, and it can also negatively regulate retinoic acid biosynthesis by diverting retinol away from oxidative activation [42]. Therefore, we speculate higher level of retinyl ester found in the female worms from SCID mice logically means lower level of lecithin, which should be consumed to synthesize retinyl ester. What was consistent with this inference was that PC(22:6/20:1) (HMDB0008735, with full name as 1-docosahexaenoyl-2-eicosenoyl-sn-glycero-3-phosphocholine, or traditional name as lecithin) happened to be decreased in the female worms from SCID mice when compared with those from BALB/c mice in the results (Figure 5). So, we speculated that the accumulated retinyl ester in the female worms from SCID mice resulted in insufficient formation of the active retinoic acid. It is known that retinoic acid is the meiosis-inducing factor in both sexes, and inhibition of retinoic acid biosynthesis would markedly suppresses gametogenesis [37-41, 43-54]. In addition, overexpression of LRAT will favor retinyl ester formation, which would disrupt retinol homeostasis and interrupt the ability of downstream metabolites to regulate transcription of genes involved in various biological processes. Various cancer cells have been found to have low levels of LRAT and retinyl ester levels. Overexpression of LRAT or increased level of retinyl esters themselves makes cells more sensitive to carcinogen-induced tumorigenesis and leison [55, 56]. So, insufficient retinoic acid biosynthesis due to prevailing retinyl ester formation could be a significant cause for the retarded development and declined fertility appeared in female worms from SCID mice in this study. Meanwhile, lecithin is a source of several active compounds: choline and its metabolites are needed for several physiological purposes, including cell membrane signaling and cholinergic neurotransmission during growth and reproduction [57, 58]. So, excessive lecithin consumption in retinyl ester formation probably lead to worse effect in the development and reproduction of schistosomes besides insufficient retinoic acid biosynthesis. In contrast, however, an extremely higher level of lecithin was detected in the male worms from SCID mice than BALB/c mice, with a relative fold as high as 17.53 for IS-MALE *vs.* IB-MALE. It is known that the schistosomes are dioecious trematodes, and embracing with the male worm by residing in the male’s gynaecophoric channel is crucial for the female worm to grow and sexually mature [59]. So, we speculated that insufficient interaction between the male and female worms, which manifested as decreased percent of worm pairs reported in our previous research [12], resulted in insufficient material exchange or transfer between them, such as the probable lecithin transfer from male worms to female worms. Similar alterations in fatty acyls, glycerophospholipids, purine nucleosides, imidazopyrimidines and indoles and derivatives were detected in both male and female worms from SCID mice when compared with BALB/c mice, but opposite alterations were detected in carboxylic acids and derivatives (Figure 4-A, C, E). Common decrease in glycerophospholipids synthesis with lysoPCs, lysoPEs and PCs as the top three alter metabolites, one of the main functions of which is to serve as a structural component of biological membranes [60, 61], indicated attenuated parasite establishment with smaller body size and attenuated reproduction due to potentially deficient glycerophospholipids in worms from SCID mice. Sphingolipids are commonly believed to protect the cell surface against harmful environmental factors [62, 63], and arachidonic acid is involved in cellular signaling as a lipid second messenger as a polyunsaturated fatty acid present in the phospholipids [64, 65]. Their involved metabolic pathways of important biological significance such as sphingolipid metabolism and arachidonic acid metabolism were also found abnormal with decreased sphingolipids in male and arachidonic acid in female worms from SCID mice, respectively, which may be also associated with the developmentally stunted worms.

**Figure 5.**
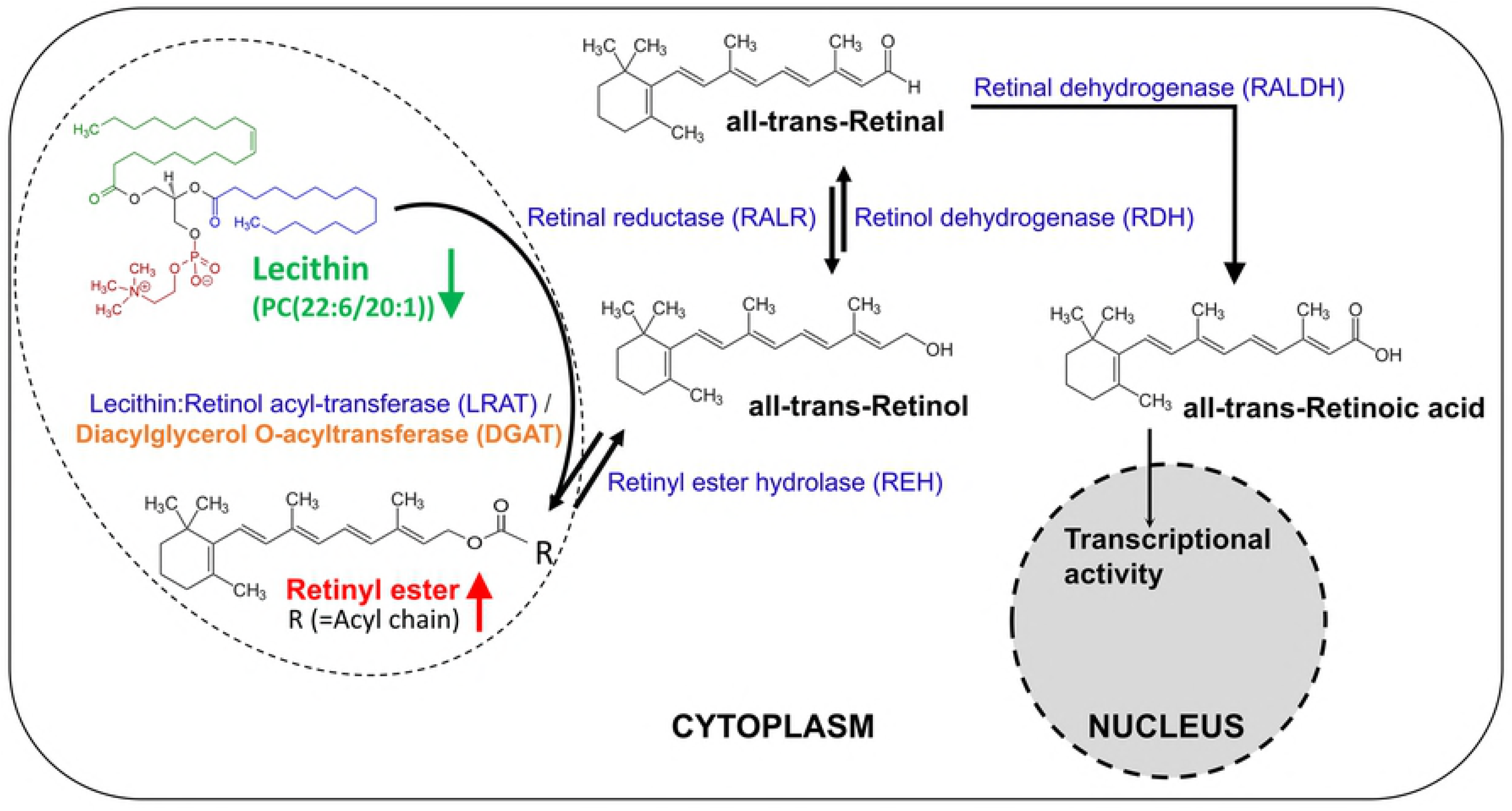
The metabolism of retinoids. Retinal can be originally formed by the cleavage of proretinoid carotenoids such as beta-carotene by the enzyme BCMO1 (not shown here). Retinol is formed by the reversible reduction of retinal by one of the retinal reductase family members. The enzyme lecithin:retinol acyl-transferase (LRAT) (with an orthologue gene called diacylglycerol O-acyltransferase (DGAT) in *Schistosoma japonicum)* synthesizes retinyl esters by transferring a fatty acyl moiety from the sn-1 position of membrane phosphatidyl choline such as lecithin (PC(22:6/20:1)) to retinol. Unesterified retinol is liberated from retinyl ester stores by the catalysis of a retinyl ester hydrolase (REH). Retinol is oxidized by catalysis of one retinol dehydrogenase (RDH) to retinal, which is then irreversibly oxidized by one retinal dehydrogenase (RALDH) to form transcriptionally active retinoic acid. Retinoic acid is finally oxidized/catabolized to more water-soluble hydroxy- and oxo- forms by one of several cytochrome P450 enzyme family members.

Furthermore, the level of tryptophan, an essential amino acid, was found decreased in both male and female worms from SCID mice. Tryptophan acts as a precursor for the synthesis of the neurotransmitters melatonin and serotonin and then, any reduction in tryptophan will lead to a number of conditions or diseases e.g. dermatitis and psychiatric symptom – depression in animals, it seems in this study to contribute to inhibiting of the growth and reproduction of worms finally [66-71]. Ergothioneine is a product of plant origin that accumulates in animal tissues and a naturally occurring metabolite of histidine that has antioxidant properties though its physiological role in vivo is undetermined [72]. Decrease of ergothioneine was found in female worms from SCID mice, which indicated increased susceptibility to oxidative damage in them [73-75].

In conclusion, the identified differential metabolites and their involved metabolic pathways are likely associated with the abnormalities in growth and development of *S. japonicum* worms in SCID mice when compared with those in BALB/c mice. Differential alterations in metabolic profiles between male and female worms from SCID mice when compared with BALB/c mice indicated the degree and mechanism of the influence the host on male and female worms were different. Our data has demonstrated the great ability of LC-MS/MS-based metabolomics to detect a broad range of differential metabolites in worms that strongly distinguished between their different hosts - the SCID mice and BALB/c mice. The above mentioned differential metabolites, together with the others not mentioned here in particular, need further verification and investigations for their underlying mechanisms in the regulation of growth and development of schistosome. This will, as a result, greatly facilitate the discovery of new drugs and vaccines against schistosome and schistosomiasis.

## Supporting Information

**Figure S1.**
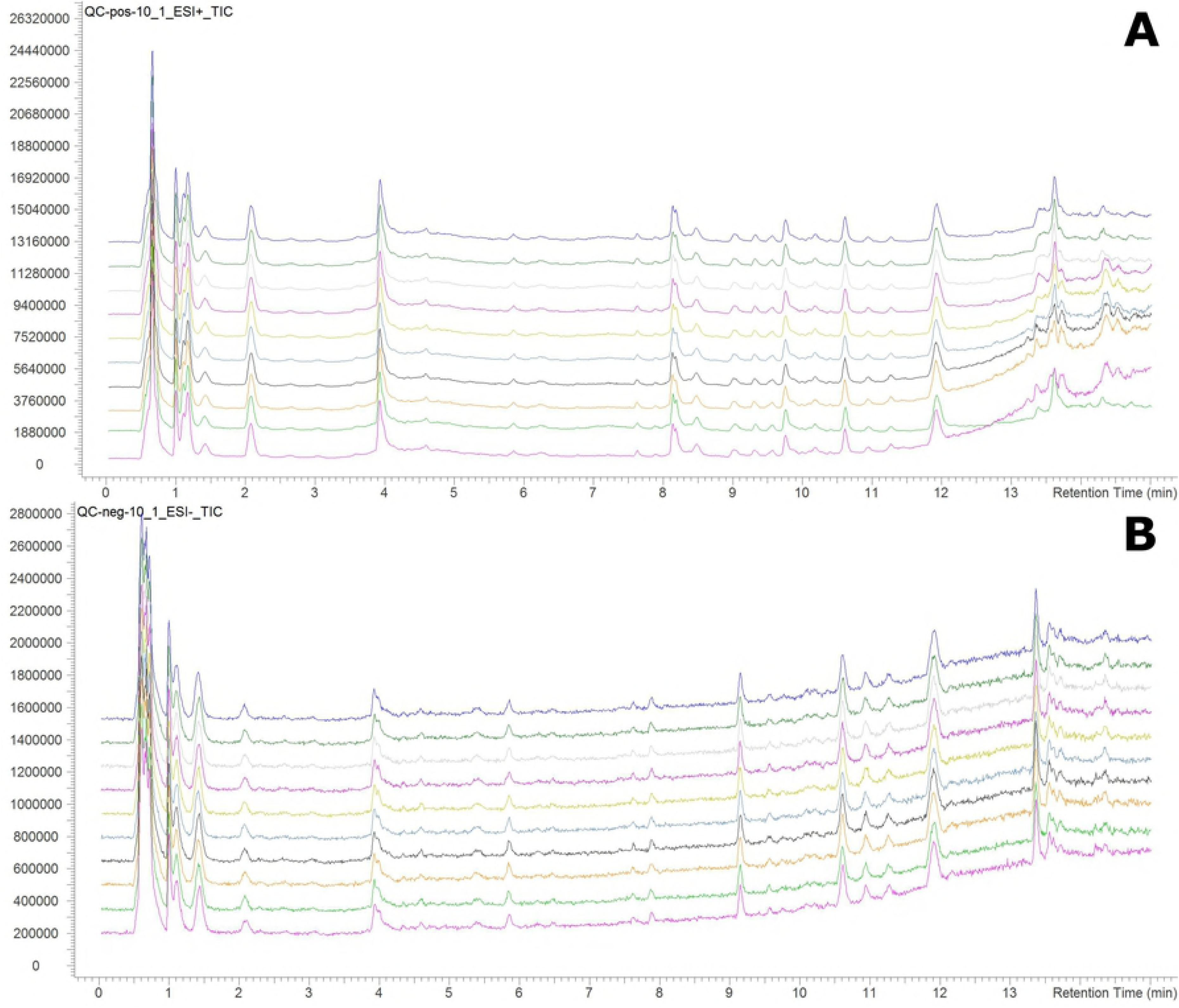
The stacked total ion chromatograms of QC sample. **A**: The stacked total ion chromatogram of QC sample in ESI+ mode. **B**: The stacked total ion chromatogram of QC sample in ESI-mode. The bars on x-axis represent the retention time (0∼15 min) and the bars on y-axis represent the total ion strength.

**Figure S2.**
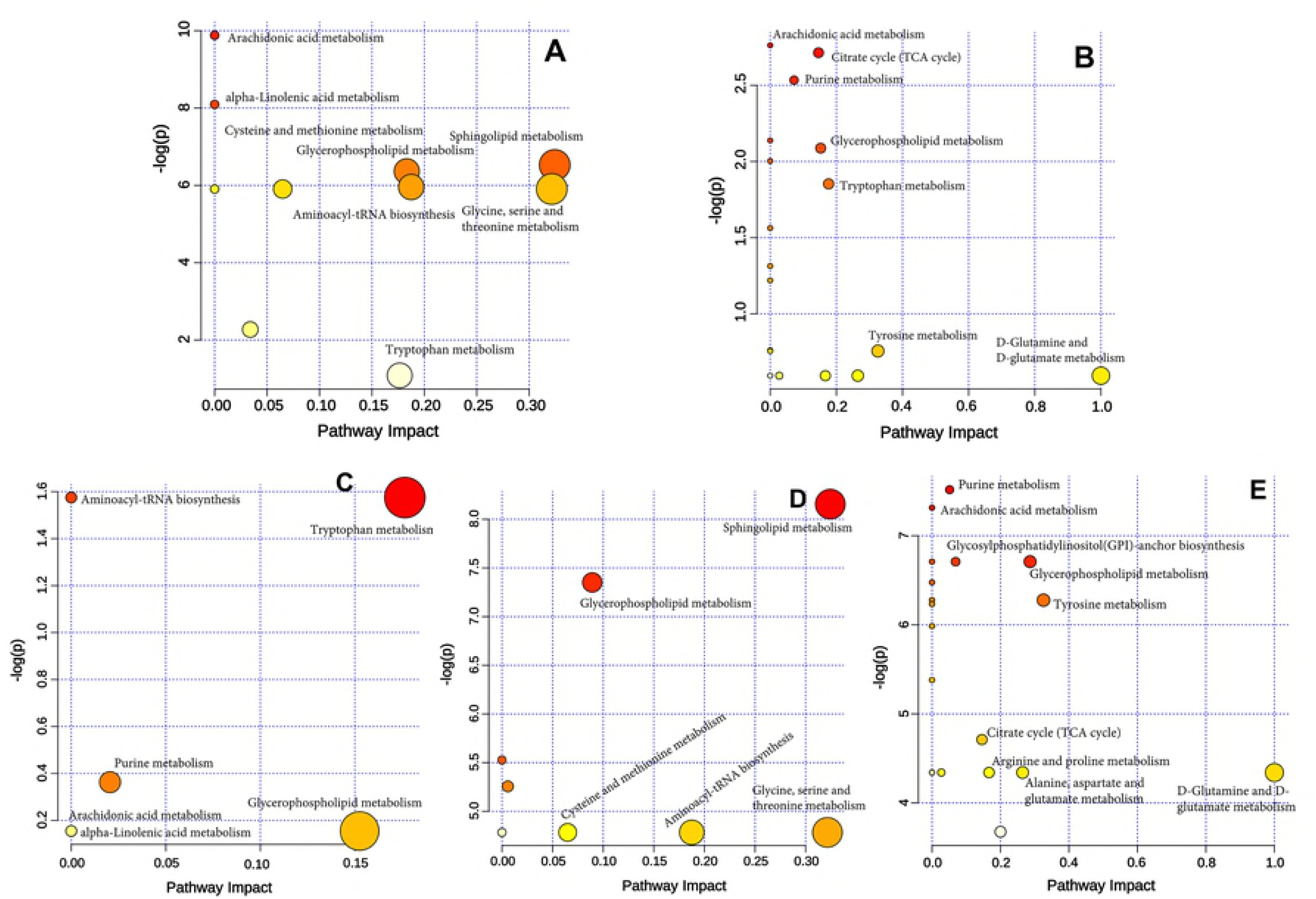
Summary of the aberrant metabolic pathways based on the significantly altered metabolites of the comparison groups as analyzed by Pathway Analysis of *MetaboAnalyst*. Plots show the matched pathways depicted according to *P*-value from pathway enrichment analysis and pathway impact score from pathway topology analysis. **A**: Enriched metabolic pathways based on the differential metabolites between IS-MALE *vs.* IB-MALE. **B**: Enriched metabolic pathways based on the differential serum metabolites between IS-FEMALE *vs.* IB-FEMALE. **C**: Enriched metabolic pathways based on the differential metabolites common in IS-MALE *vs.* IB-MALE and IS-FEMALE *vs.* IB-FEMALE. **D**: Enriched metabolic pathways based on the differential metabolites distinct in IS-MALE *vs.* IB-MALE. **E**: Enriched metabolic pathways based on the differential metabolites distinct in IS-FEMALE *vs.* IB-FEMALE. Color gradient and circle size indicate the significance of the pathway ranked by *P*-value (yellow: higher *P*-values and red: lower *P*-values) and pathway impact score (the larger the circle the higher the impact score), respectively. Significantly affected pathways with low *P*-value and high pathway impact score are identified by name.

### Supporting table legends

**Table S1. Differential metabolites of male worms from SCID mice compared with those from BALB/c mice.** This table contains a list of the differential metabolites between male worms from SCID mice and male worms from BALB/c mice.

(DOCX)

**Table S2. Differential metabolites of female worms from SCID mice compared with those from BALB/c mice.** This table contains a list of the differential metabolites between female worms from SCID mice and female worms from BALB/c mice.

(DOCX)

**Table S3. Metabolite set enrichment of the differential metabolites of male worms between SCID mice and BALB/c mice.** Metabolite sets are enriched of the differential metabolites between male worms from SCID mice and male worms from BALB/c mice.

(DOCX)

**Table S4. Metabolite set enrichment of the differential metabolites of female worms between SCID mice and BALB/c mice.** Metabolite sets are enriched of the differential metabolites between female worms from SCID mice and female worms from BALB/c mice.

(DOCX)

**Table S5. Metabolite set enrichment of the common differential metabolites in male and female worms from SCID mice.** Metabolite sets are enriched of the common differential metabolites between male worms and female worms from SCID mice compared with those from BALB/c mice.

(DOCX)

**Table S6. Metabolite set enrichment of the male worm-specific differential metabolites between SCID mice and BALB/c mice.** Metabolite sets are enriched of the male worm-specific differential metabolites between SCID mice and BALB/c mice.

(DOCX)

**Table S7. Metabolite set enrichment of the female worm-specific differential metabolites of those between SCID mice and BALB/c mice.** Metabolite sets are enriched of the female worm-specific differential metabolites between SCID mice and BALB/c mice.

(DOCX)

**Table S8. Pathway analysis of the differential metabolites of male worms between SCID mice and BALB/c mice.** This list contains the enriched metabolic pathways of the differential metabolites of male worms between SCID mice and BALB/c mice.

(DOCX)

**Table S9. Pathway analysis of the differential metabolites of female worms between SCID mice and BALB/c mice.** This list contains the enriched metabolic pathways of the differential metabolites of female worms between SCID mice and BALB/c mice.

(DOCX)

**Table S10. Pathway analysis of the common differential metabolites in male and female worms from SCID mice.** This list contains the enriched metabolic pathways of the common differential metabolites between male worms and female worms from SCID mice compared with those from BALB/c mice.

(DOCX)

**Table S11. Pathway analysis of the male worm-specific differential metabolites between SCID mice and BALB/c mice.** This list contains the enriched metabolic pathways of the male worm-specific differential metabolites between SCID mice and BALB/c mice.

(DOCX)

**Table S12. Pathway analysis of the female worm-specific differential metabolites between SCID mice and BALB/c mice.** This list contains the enriched metabolic pathways of the female worm-specific differential metabolites between SCID mice and BALB/c mice.

(DOCX)

